# Don’t Leave the Past Behind: How Larval Experience Shapes Pupal Antipredator Response in *Aedes aegypti*

**DOI:** 10.1101/2024.02.02.578532

**Authors:** Kanika Rawat, Akshaye Anand Bhambore, Kavita Isvaran

## Abstract

Animals use predation encounters or risk experiences to influence their future antipredator responses. Such carryover effects of predation can benefit them by enhancing their antipredator behaviour and thereby decreasing their risk of mortality through predation. Despite these fitness benefits, behavioural carryovers of predation past may not be a common phenomenon in complex life cycles. Complex life cycles pose a unique evolutionary and physiological challenge to behavioural carryovers since every life stage is distinct in morphology, physiology, and function. Each life stage of a complex life cycle is expected to evolve its independent response based on the real-time threat level and manage the trade-offs accordingly. Along with the evolutionary challenge, physiological barriers can hamper behavioural carryovers, especially in holometabolous insects, where we observe extensive tissue remodelling and developmental compartmentalisation. We investigated behavioural carryover in the holometabolous mosquito model system, *Aedes aegypti*. We asked whether predation risk during a life stage carries over to the subsequent stage, influencing its behaviour, or if the next life stage responds according to its threat environment. *Aedes aegypti* has four major stages– egg, larva, pupa, and adult. We examined the effect of predation-risk experience across larval and pupal stages. Larval and pupal stages differ in morphology, physiology and function. They share the same habitat and, therefore, similar threats. We manipulated the threat of predation experienced by larvae and investigated its influence on pupal behaviour. We found behavioural carryover in the pupal stage for the first time and discovered exciting interactions between past experiences and the current threat environment. Our study underscores the crucial role of predation pressure in shaping the evolution of complex life cycles, emphasising the significance of early experiences with predators in influencing behavioural traits across distinct life stages.

## Introduction

The high cost of predation has led to the evolution of diverse antipredator strategies that animals deploy either to avoid predator attacks– such as camouflage, mimicry, aposematism; or to fight and escape the attacks– use of spines and armours, chemical defences (Breed & Moore, 2016; Caro, 2005; Sugiura, 2020). While some of these strategies are relatively fixed in expression, others are more flexible during an individual’s lifetime (as explained by Sergio et al., 2021). An essential class of flexible antipredator strategies are behavioural tactics. Animals use behavioural responses such as freeze, fight and flight to survive encounters with their predators (Breed & Moore, 2016; Denver, 2010); however, many more behavioural decisions are made to avoid the encounters in the first place (Lima & Dill, 1990). Behaviours such as vigilance, socialising, and seeking refuge decrease the probability of predator attacks. The time and energy spent on these behaviours are traded against crucial behaviours such as foraging and mating (Dill, 1987; Lima & Dill, 1990). Therefore, we see animals optimising their behaviour as per their vulnerability to a potential predator to improve their chances of survival.

To assess the risk in their surroundings, animals use cues such as illumination, microhabitat structure, and auditory, olfactory and physical cues of predators (Nersesian et al., 2012; Thorson et al., 1998). For example, white-footed mice decrease their space use during full moon nights when they hear predator calls (A. Schmidt, 2006). Apart from using information from their immediate environment, animals can also use their past experiences to inform their expectation of risk and modulate their current antipredator behaviour. For example, snakes that are captured before are more likely to flee during the second capture (Gregory, 2013), zebrafish react faster to a predatory stimulus as they accumulate experience (Dill, 1974), and female Greater rheas trained with visual predator cues reduce their movement in the wild (Cortez et al., 2018).

Such behavioural carryovers of predation risk have also been reported being advantageous between generations (Keiser & Mondor, 2013; Shine & Downes, 1999; Storm & Lima, 2010) and from eggs/embryos to post-hatchings (Crowder & Ward, 2022; Garcia et al., 2017; Mathis et al., 2008). For example, offspring of two-spotted spider mites with predation experience prefer residing on the leaf blade rather than near the leaf vein, reducing the probability of predator encounters (Freinschlag & Schausberger, 2016). Exposure to predation cues during the embryonic stage elicits an antipredator response by the Indian skipper frog tadpoles in the presence of predator kairomones (Supekar & Gramapurohit, 2024). While behavioural carryovers from predation past are expected to be advantageous and prevalent, as shown in the review by Tariel et al., 2020, the complexity of a life cycle may pose challenges to their evolution.

A complex life cycle refers to a life history where an organism switches its morphology, physiology and function at different life stages associated with different habitats (Wilbur, 1980). Due to these differences, life stages are expected to evolve independently to adapt to their environment. Such a life history can limit behavioural carryover due to evolutionary and physiological constraints (Moran, 1994). Although there is some evidence in favour of behavioural carryover between post-birth life stages in complex life cycles of fish and amphibians (e.g., Garcia et al., 2019; Lönnstedt et al., 2012), life cycles with extreme metamorphosis are still virtually unexplored.

Life cycles of holometabolous insects exemplify complete metamorphosis (Gilbert, 2000). Unlike amphibians and marine fish, they experience extreme tissue remodelling and compartmentalisation during metamorphosis (Moran, 1994; Rolff et al., 2019). Such life cycles go through egg-larva-pupa stages to reach the adult stage. We see evidence for the independent evolution of some traits between life stages (larval and adult) in holometabolous insects (Collet et al., 2023; League et al., 2017). However, we must determine whether this independent evolution also decouples risk assessment and hinders behavioural carryover of early predation-risk experience across life stages.

One recent study has looked at the behavioural carryover of predation risk in a holometabolous life cycle. Krama et al., 2023 showed that the *D. melanogaster* adults with larval experience of predation risk show a behaviour change and survive better in the presence of predators than naive adults. Studies on antipredator defence and the carryover effects of predation often neglect the pupal stage (Lindstedt et al., 2019), perhaps because of the developmental role of the pupal stage relative to the larval (feeding) and adult (reproductive) stages. Larva accumulates energy for transformation into an adult through the pupal stage. The pupal stage is usually immobile. It does not acquire energy and relies on the larval accumulation of resources to survive and transform into an adult. As much as these traits make a pupal study less engaging, they make pupa a crucial and sensitive link between the larval and adult stages as it has to manage trade-offs with limited energy under the constant threat of predation. Moreover, pupae in many taxa can show behavioural defences that need more exploration (Lindstedt et al., 2019).

To understand how past predation-risk experience affects the behaviour of the subsequent stage in a complex life cycle, we aim to focus on the behavioural carryover of larval experience onto the pupal stage in *the Aedes aegypti* mosquito. *Aedes aegypti* pupa is a motile, tissue remodelling stage that relies on the feeding larval stage for energy (Nelson, 1986). *Aedes aegypti* larvae and pupae are found in natural ephemeral pools and artificial water containers (Dharmamuthuraja et al., 2023; Rao et al., 1970). The larval stage lasts six to eight days, and the pupal stage lasts two to three days before metamorphosing into an adult (Christophers, 1960). Since both the aquatic stages share the same habitat, they experience similar predation-risk conditions (Kumar & Hwang, 2006; Vinogradov et al., 2022). Larvae employ tactics such as reducing movement and changing habitat use to safeguard themselves from predators (Chandrasegaran et al., 2018; Sih, 1986). *Aedes aegypti’s* motile pupae can also sense threats and protect themselves using different behavioural traits.

Pupae rest and breathe at the surface (Nelson, 1986). They tend to dive whenever the water is disturbed, and shadows are cast on the water surface (Romoser & Lucas, 1999). The diving behaviour has been investigated in *Culex pipiens* pupae, actively chased by guppy fish, by Awasthi et al., 2012. Recently, Chandrasegaran et al., 2020 studied pupal-antipredator response in *Aedes aegypti* under constant predation risk, using chemical and visible predator cues. This study reported that pupae switch their habitat use under predation risk and visit the water surface less frequently than in the absence of the risk.

To test whether larval predation-risk experience affects pupal antipredator behaviour, we used chemical and visual cues of predators to manipulate the perception of predation risk in larval growth environments since they respond to predation cues (Chandrasegaran et al., 2018; Sih, 1986). We created two larval growth conditions: one with predation risk and one without predation risk. We then tested for carryover effects on pupal behaviour under two environments– no immediate or immediate threat. We expected pupae with larval predation-risk experience to perceive their environment as risky and modify their behaviour to reduce the likelihood of encountering a potential predator. Behaviour modification was expected to be intensified under an immediate threat environment. Alternatively, the pupal stage might only respond to real-time risk and not get affected by the larval experience if physiological and evolutionary constraints prevent the behavioural carryover.

Since mosquito larvae usually grow under high-density and competitive environments (Arrivillaga & Barrera, 2004; Barrera et al., 2006; Strickman & Kittayapong, 2003), we studied the carryover under resource-limited conditions. We also explored what happens when the resource limitation is relaxed. We expected to observe differences in activity and antipredator behaviour in pupae due to the costly nature of the behaviour (Lucas & Romoser, 2001). Therefore, we also explored whether and how carryover trends vary between resource-limited and resource-rich conditions due to the associated energetic costs.

We extensively observed the designated escape response– diving behaviour and recorded space use and overall activity patterns, as animals are known to modulate these tactics to avoid predator encounters (Abramsky et al., 1996; Formanowicz & Bobka, 1989; Valeix et al., 2009; Van Buskirk & Yurewicz, 1998).

## Materials and Methods

### Predators and mosquito eggs collection

We used granite ghost dragonfly (*Bradinopyga geminata*) nymphs as the predators since they feed on *Aedes aegypti* larvae and pupae in their aquatic habitat (Sharma et al., 2020; Venkatesh & Tyagi, 2013). Granite ghost dragonflies are endemic to South Asia (Kalkman et al., 2020), and their nymphs are found in rock pools and small water tanks (Frazer, 1936). The nymphs are sit-and-wait predators that occasionally swim and chase their prey (Subramanian, 2009, personal observations). We collected granite ghost dragonfly nymphs from different ephemeral rock pools around Bengaluru during the monsoon (2021-2022). We maintained these nymphs under laboratory conditions by feeding them *Aedes aegypti* larvae and pupae and changing their container water regularly. Large-sized nymphs that measured 1.5 cm to 2.0 cm were chosen for the experiment as they show a high predation rate and no preference for different larval instars (Shad & Andrew, 2013). We collected *Aedes aegypti* eggs from our mosquito colony at the Indian Institute of Science, Bengaluru (set up in 2013, eggs sourced from the National Malaria Research Institute). The mosquito colony is maintained at 12:12 hours of light:dark cycle, 70-80% relative humidity and 25°C-28°C.

### Experimental setup

#### Larval growth phase

We hatched *Aedes aegypti* eggs in tap water. After 24 hours of hatching, we collected the first instar larvae for our experiment. Half of the first instar population was grown under predation-risk conditions (’experienced’ treatment), and the other half was grown under no predation-risk conditions (’naive’ treatment). We counted 45 first instar larvae and added them to brown plastic trays (25 cm diameter and 5 cm height) with two litres of tap water and larval food. Larval food was prepared by grinding and mixing dog biscuits (Glenand Dog Biscuits) and yeast (in a 3:2 ratio). We placed two dragonfly nymphs, caged in 50 ml plastic containers, at the centre of the experienced treatment trays to provide visual cues to the larvae. In contrast, we placed two 50 ml plastic containers without nymphs in the naive treatment trays. We maintained twelve such trays in one experiment block, with six replicates per treatment. We performed seven blocks of the experiment with two different food conditions: resource-limited (0.15 g) and resource-rich (3.0 g) from January to December 2022. The food levels and conspecific density were chosen based on the lab protocols (Sharma et al., 2020) and preliminary experiments testing larval growth across a range of food levels. 0.15 g food level provides a competitive environment, typically experienced by mosquito larvae, while 0.3 g relaxes the resource limitation.

Two blocks were successfully completed under resource-limited conditions, while three blocks were completed under resource-rich conditions. Throughout all experiment blocks, we maintained a 12:12 hours light:dark cycle, 60%-75% relative humidity, and a temperature range of 24°C-27°C.

We fed the experiment nymphs with six larvae (third and fourth instars) and pupae per day from our mosquito colony during the seven to eight days of the larval growth period. The nymph containers were decanted into the trays and filled again using the same tray water. We did this twice daily (around 9 am and 9 pm) to add chemical cues (dead remains of immature mosquitoes and nymphs’ faecal matter) of predation to the experienced treatment trays. After the larvae reached a sufficiently large size (fourth instar), we replaced the nymphs’ plastic containers with perforated containers to ensure a continuous dispersal of predation cues. We maintained a uniform handling protocol for experienced and naive treatment to account for the disturbance caused to growing larvae. The tray water level was replenished throughout the larval growth phase.

Towards the end of the larval stage, specifically on the sixth and seventh day after hatching, we observed the larvae at six-hour intervals for pupation. Once we spotted a few pupae in a tray, we discarded them as they were exposed to larval environment cues. We transferred the remaining soon-to-pupate larval population to a corresponding tray (’transfer’ tray). Transfer trays were prepared along with the larval growth phase trays to mimic the growth phase conditions sans the predation cues to house the pupal population. The transfer trays had water, food and the same density of larvae as the treatment trays to mimic conspecific cues. We removed the larvae from these transfer trays just before the transfer of soon-to-pupate treatment larvae. This transfer was crucial to maintain the naivete of the pupal stage as we were interested in looking at the behavioural carryover of larval predation-risk experience.

### Behavioural assay

We recorded pupal behaviour 12 to 36 hours after the transfer. To study the behavioural carryover, we examined pupal behaviour in the presence (predation-cue water) and the absence of immediate threat (control water) during the 12 hours of light conditions. We prepared the predation-cue water from the 1st day of the larval growth phase, where we set up tubs with six litres of tap water and maintained six nymphs per tub. These nymphs were housed in perforated containers and fed six larvae (third & fourth instars) and pupae from the colony every day until the day of the behavioural assay. The predation cues dispersed through the containers to the tub water. Control water was prepared similarly by placing perforated containers without nymphs in six litres of tap water.

On the day of recording, we transferred individual pupae to transparent plastic tanks (10 x 10 x 15 cm) filled with control or cue water till 10 cm of the height. The cue water tanks also contained one dragonfly nymph in a glass beaker for visual cues. Control tanks had just empty glass beakers. We marked the tanks at every 2 cm along the height. These markings were used to measure the behavioural traits of our interest.

We recorded the pupal behaviour in the tanks using a Sony AX43 Handycam at 4k video quality (25 frames per second) against a white background. Video recording lasted 10 minutes for each trial. We allowed pupae to acclimate for 3 minutes and recorded their behaviour in the following 7 minutes. Each pupa was recorded only once, either in control or cue water. Nymphs were not repeated in the behavioural trials. We simultaneously recorded all four combinations of larval experience and behavioural assay (Fig. 1) and randomised them spatially.

**Fig. 1:**
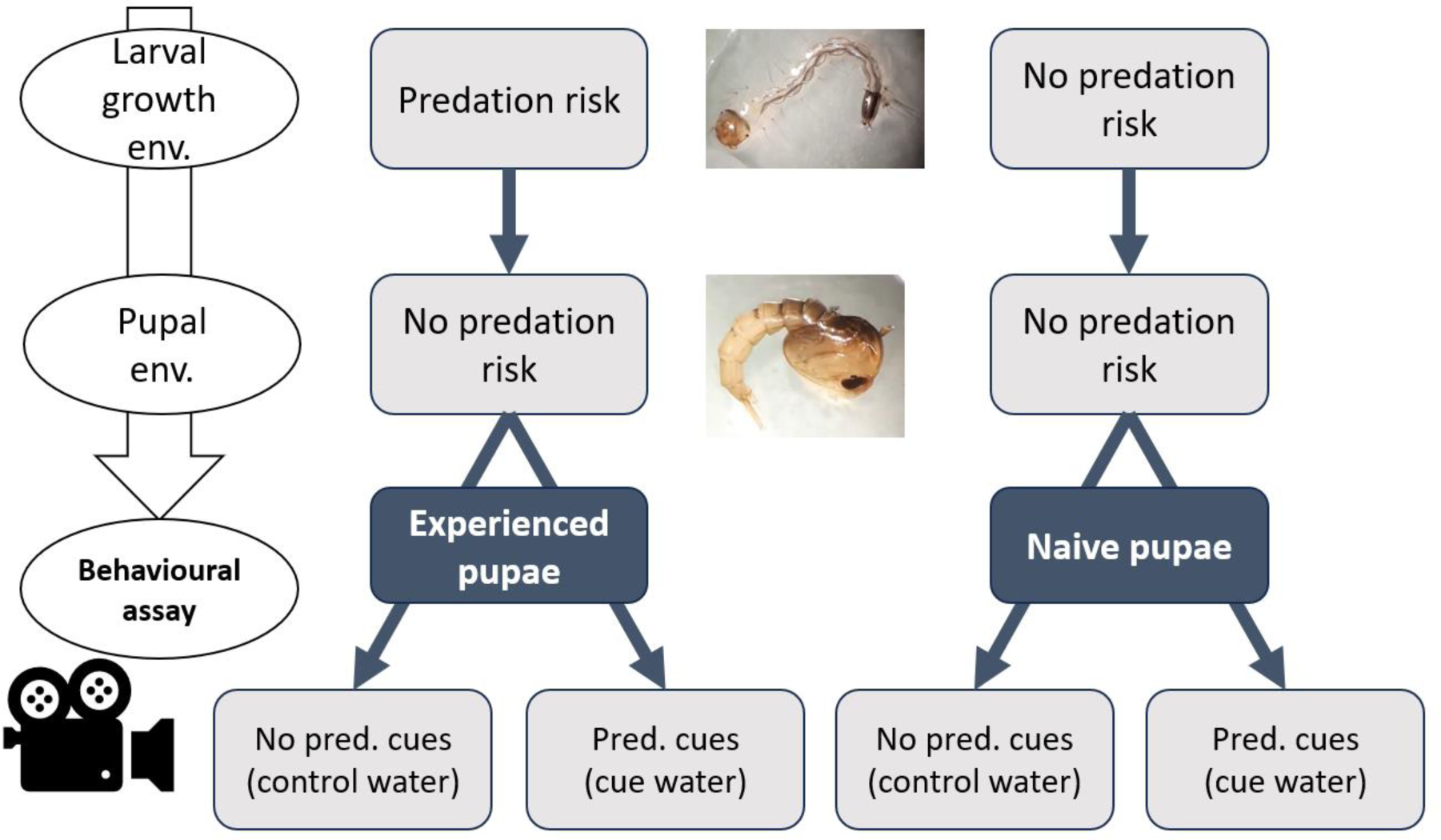
Schematic representation of the experiment design. We grew larvae in two conditions– predation risk (experienced) and no predation risk (naive). The pupal stage did not experience any predation risk in both treatments. We recorded the behaviour of experienced and naive pupae in the absence of an immediate threat (control water) and the presence of an immediate threat (predation-cue water).

We measured different behavioural traits, commonly shown by pupae, that we expected to change under predation threat– 1. Dive, 2. Space use, 3. Activity.

1. To extract dive data, we recorded all the vertical movements where the pupae broke away from the surface (Houlihan, 1971) and entered the water column. We looked at the frequency of dives in the last 7 min and the vertical depth of each dive. Vertical depth was recorded as the deepest point a pupa reaches before returning to the surface. We used the markings on the tanks to measure the dive depths at 2 cm resolution. Pupae are known to dive under threat. This behaviour can save them from predators and from getting washed away during rains (Rodríguez-Prieto et al., 2006; Romoser & Lucas, 1999; Romoser, 1975).
2. For space use data, we broadly divided the water column into the water surface, middle (top 8 cm) and bottom (last 2 cm) layers. Pupae were recorded every 10 seconds at these different locations. We used this data to look at the proportion of time spent by a pupa along the height of the water column.
3. We recorded the proportion of time spent being active by the pupa. Activity refers to visible jerky muscular contractions by pupae. For ease of data extraction, we recorded the passive behaviour data and subtracted the proportion of time spent in the passive state from 1. We recorded passive bouts of ≥ 2 seconds where pupae showed resting or floating behaviour without muscular contractions.

All the behavioural recordings were performed during the light phase in one to two days since the pupal stage lasts approximately two to three days. Due to these life cycle and experimental constraints, different pupae were at different development phases during the recordings. This can add some noise but should not affect the comparison between experienced and naive pupae. We manually extracted data from 84 individual recordings in resource-limited and 122 recordings in resource-rich conditions. Every video data was cross checked by two observers.

### Statistical analysis

Statistical analyses were performed in R version 4.2.3. We used linear and generalised linear models (Table 1) with our standard predictor variables—larval experience, behavioural assay environment, and experiment blocks—to detect their influence on various behavioural traits. The experiment block was used as a fixed effect to account for block-specific microenvironments and egg batches. We also introduced an interaction term between larval experience and the behavioural assay environment to test our prediction that the difference in the response of naive and experienced pupae should intensify under immediate threat compared to in the absence of threat.

**Table 1:** Statistical models used for multiple behavioural traits along with the rationales based on the nature of the response variables.

| Behavioural trait | Statistical test | Rationale based on response variable nature |
| --- | --- | --- |
| Dive frequency | Negative binomial GLM | Over dispersed count data |
| Dive depth | Linear regression | Continuous variable |
| Variation in dive depth | Linear regression | Continuous variable |
| Space use | Multinomial regression | Multiple categorical variables in no specific order |
| Activity | Beta regression | Proportion data |

## Results

### Diving behaviour (Normal resource-limited conditions)

Multiple aspects of diving behaviour were strongly affected by prior predation experience. All three diving traits were jointly affected by larval experience and current threat environment (likelihood ratio test for interaction term– dive frequency: Df= 1, LRT = 8.893, P value= 0.002; median dive depth: Df= 1, F value= 5.366, P value= 0.022; CV dive depth: Df= 1, F value= 4.924, P value= 0.028).

In the absence of immediate threat, dive frequency differed between experienced and naive pupae (Table 2). Experienced pupae dove around 68% more frequently than naive pupae. Further, the dive frequency of experienced pupae dipped by roughly 54% under an immediate threat environment. In contrast, naive pupae did not detectably change their dive frequency in the presence versus the absence of an immediate threat. We observed similar patterns in dive depth, where experienced pupae performed shallow dives compared with naive pupae. In the absence of threat, the median dive depth for experienced pupae was roughly 4 cm, 33% less than the naive pupae, 6 cm. However, under immediate threat, the experienced pupae performed deeper dives, 7.5 cm, showing an increment of 92%, whereas naive pupae showed a weak and variable response. We also found more variation in the dive depth of experienced pupae, where SD was 67% of the mean compared to 51% for naive pupae. Under immediate threat, the SD of dive depth decreased to 53% for experienced pupae and remained around 55% for naive pupae (Fig. 2).

**Fig. 2:**
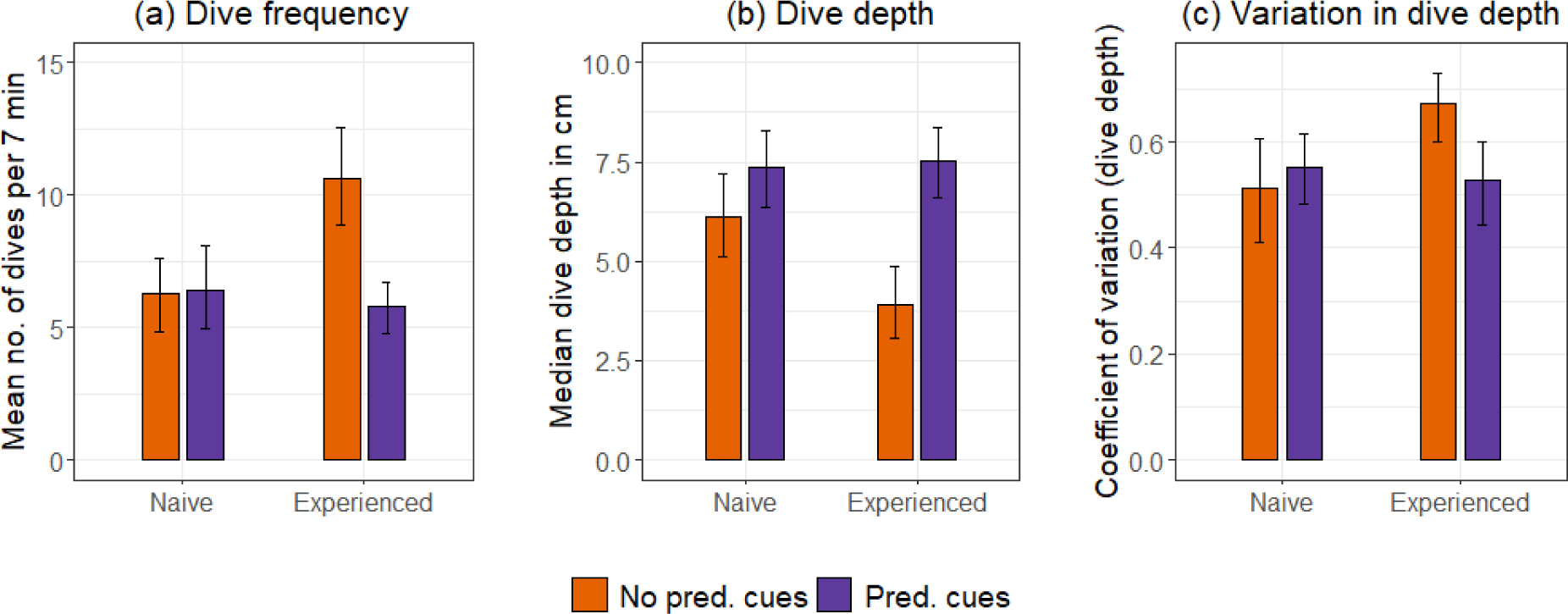
Comparison of (a) dive frequency, (b) median dive depth, and (c) coefficient of variation (CV) of dive depth between naive and experienced pupae at two different threat levels under normal resource-limited conditions. Orange represents the absence (no predation cues), and violet represents the presence of an immediate threat (predation cues). The means with 95% bootstrap confidence intervals are shown.

**Table 2:**
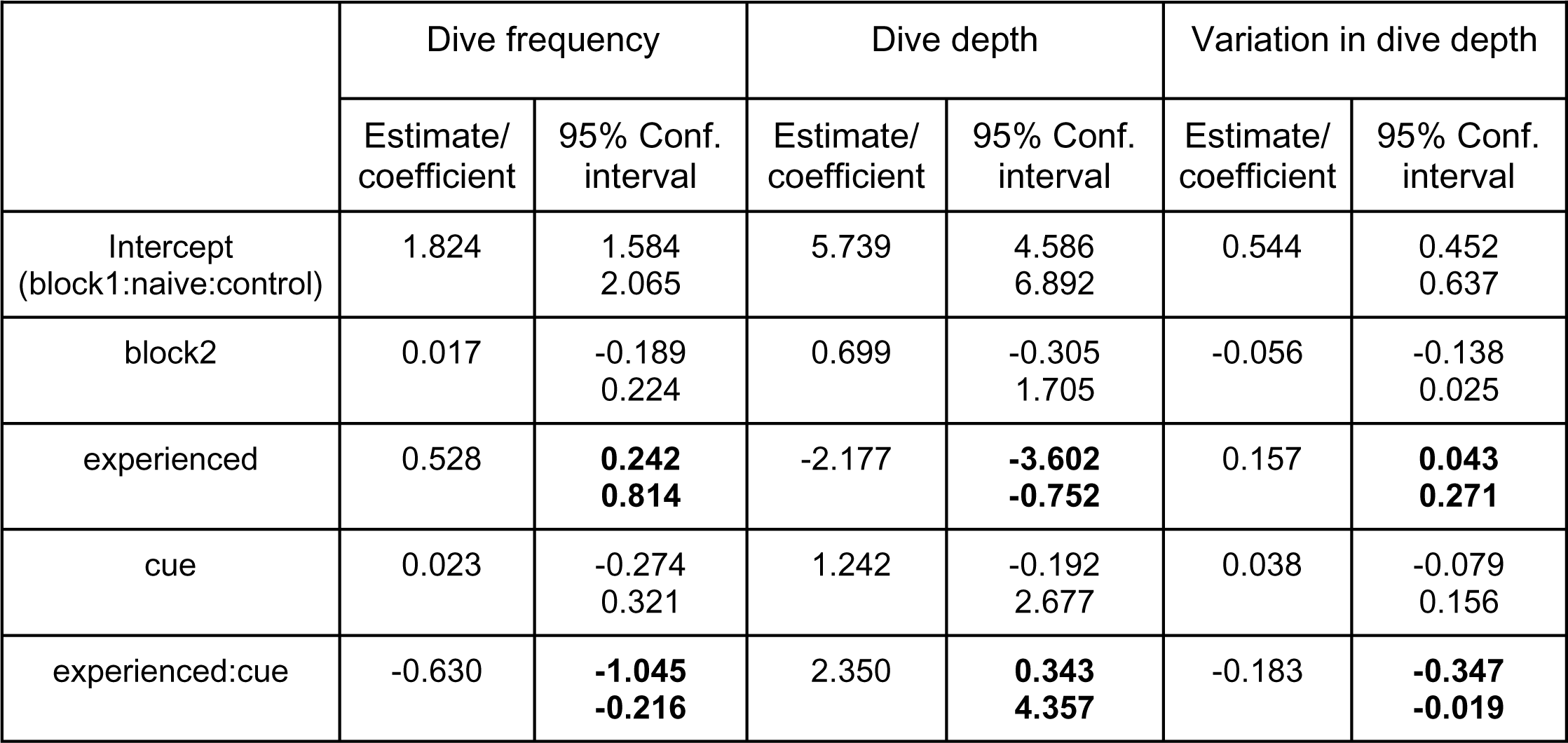
Model estimates with 95% confidence intervals from three models: negative binomial GLM with dive frequency and linear models with median dive depth and CV dive depth as response variables. We estimated the effects of the larval growth environment (experienced and naive), behavioural assay environment (predation cue and control water) and their interaction on the response variables under typical resource-limited conditions. Entries in bold indicate that the effects are detectable.

### Space use and Activity (Resource-limited conditions)

Prior experience did not detectably affect space use patterns; immediate threat did. Both experienced and naive pupae moved away from the middle column in the immediate threat environment (Table 3 & Table 4). Pupae, irrespective of their larval experience and immediate threat level, preferred spending time at the surface. Pupae spent roughly 50%, 35% and 15% of their time at the surface, middle column and the bottom layer, respectively. Similarly, pupae spent approximately similar time being active as passive, irrespective of the larval predation-risk experience and immediate threat environment (Fig. 3).

**Fig. 3:**
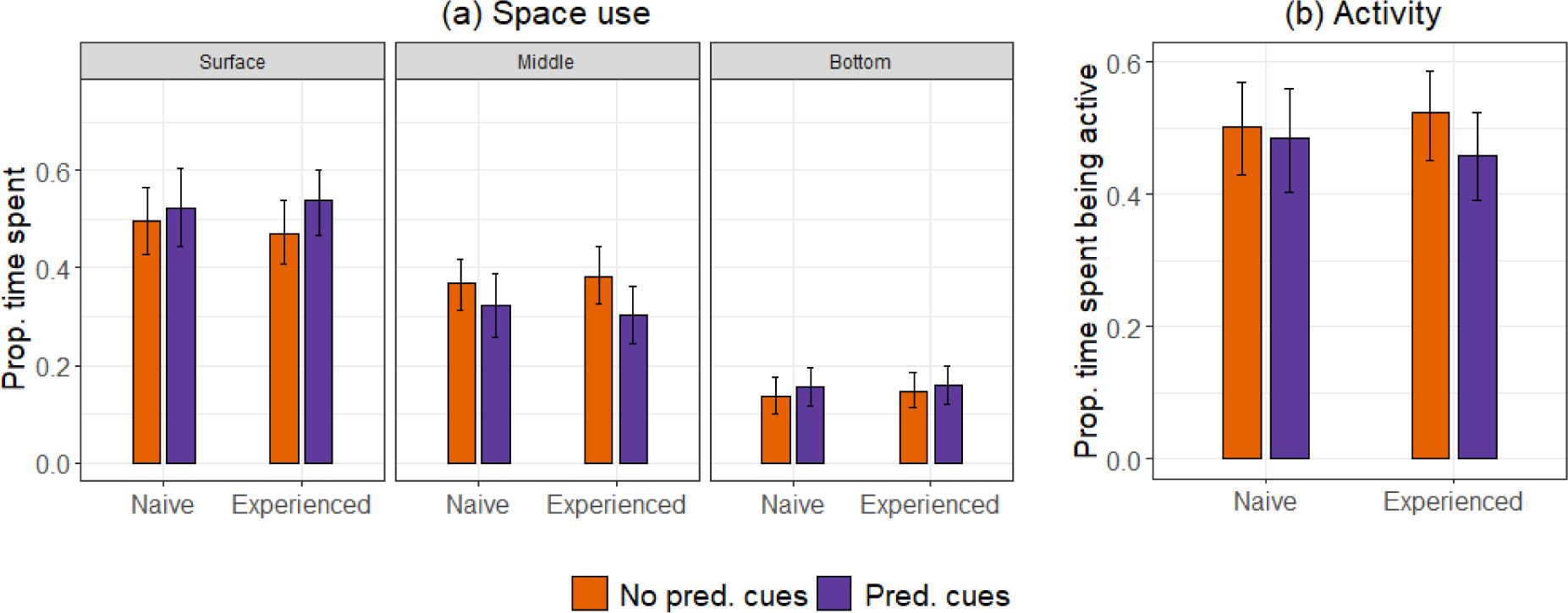
(a) The plots compare the proportion of time spent at the surface, middle and bottom layers by naive and experienced pupae across two threat levels (no predation cues and predation cues) under resource-limited conditions. (b) Comparison of the proportion of time spent being active by experienced and naive pupae across two threat levels. The means with 95% bootstrap confidence intervals are shown.

**Table 3:** Model estimates with 95% confidence intervals from two models: a multinomial model with proportions of time spent at the surface and bottom layers relative to the middle layer as response variables; a beta regression GLM with ‘proportion time spent being active’ as the response variable. We estimated the effects of the larval growth environment (experienced and naive), behavioural assay environment (predation cue and control water), and their interaction on the response variables under resource-limited conditions. Entries in bold indicate that the effects are detectable.

|  | Space use:<br>Surface/Middle<br>column |  | Space use:<br>Bottom layer/Middle<br>column |  | Activity |  |
| --- | --- | --- | --- | --- | --- | --- |
|  | Estimate/<br>coefficient | 95% Conf.<br>interval | Estimate/<br>coefficient | 95% Conf.<br>interval | Estimate/<br>coefficient | 95% Conf.<br>interval |
| Intercept<br>(block1:naive:control) | 0.456 | 0.151<br>0.761 | -0.978 | -1.423<br>-0.534 | -0.401 | -0.760<br>-0.043 |
| block2 | -0.297 | **-0.571**<br>**-0.023** | -0.012 | -0.396<br>0.371 | 0.385 | **0.066**<br>**0.703** |
| experienced | -0.100 | -0.480<br>0.280 | 0.029 | -0.514<br>0.573 | 0.172 | -0.276<br>0.621 |
| cue | 0.180 | -0.206<br>0.566 | 0.260 | -0.285<br>0.807 | -0.031 | -0.480<br>0.417 |
| experienced:cue | 0.193 | -0.354<br>0.740 | 0.047 | -0.718<br>0.812 | -0.085 | -0.720<br>0.548 |

**Table 4:** Likelihood ratio tests for the response variables– proportion time spent at surface, middle, and bottom layers; proportion time spent being active under resource-limited conditions. We first assessed the interaction term by comparing models with the interaction term, ‘Larval growth env:Behavioural assay env’, and the nested simple models (without the interaction term). Models with the interaction term did not explain the data better than the simpler models. We used the models without the interaction term to assess the statistical significance of the main effects (larval growth environment, behavioural assay environment) through likelihood ratio tests. We used the ‘lrtest’ command from the package, ‘lmtest’, to perform the likelihood ratio tests for space use and activity models.

|  | Space use |  |  | Activity |  |  |
| --- | --- | --- | --- | --- | --- | --- |
| Deleted parameter | Likelihood ratio (chisq) | Degrees of freedom | P value | Likelihood ratio (chisq) | Degrees of freedom | P value |
| Larval growth env:Beh assay env | 3.042 | 2 | 0.218 | 0.07 | 1 | 0.791 |
| Block | 32.069 | 2 | <0.0001 | 5.535 | 1 | 0.018 |
| Larval growth env | 0.498 | 2 | 0.779 | 0.640 | 1 | 0.423 |
| Beh assay env | 25.968 | 2 | <0.0001 | 0.206 | 1 | 0.649 |

### Diving behaviour (Resource-rich conditions)

Experienced pupae were similar to naive pupae under resource-rich conditions in most measured traits. We could detect a difference in variation in dive depths (Table 5). Experienced pupae showed higher variation in their dive depth than naive pupae under no immediate threat. While naive pupae dove with 52% variation around the mean, experienced pupae varied in their dive depths by 63% (Fig. 4). This difference was statistically detectable in the interaction model but not when assessed as a whole across both assay environments (Table 6).

**Fig. 4:**
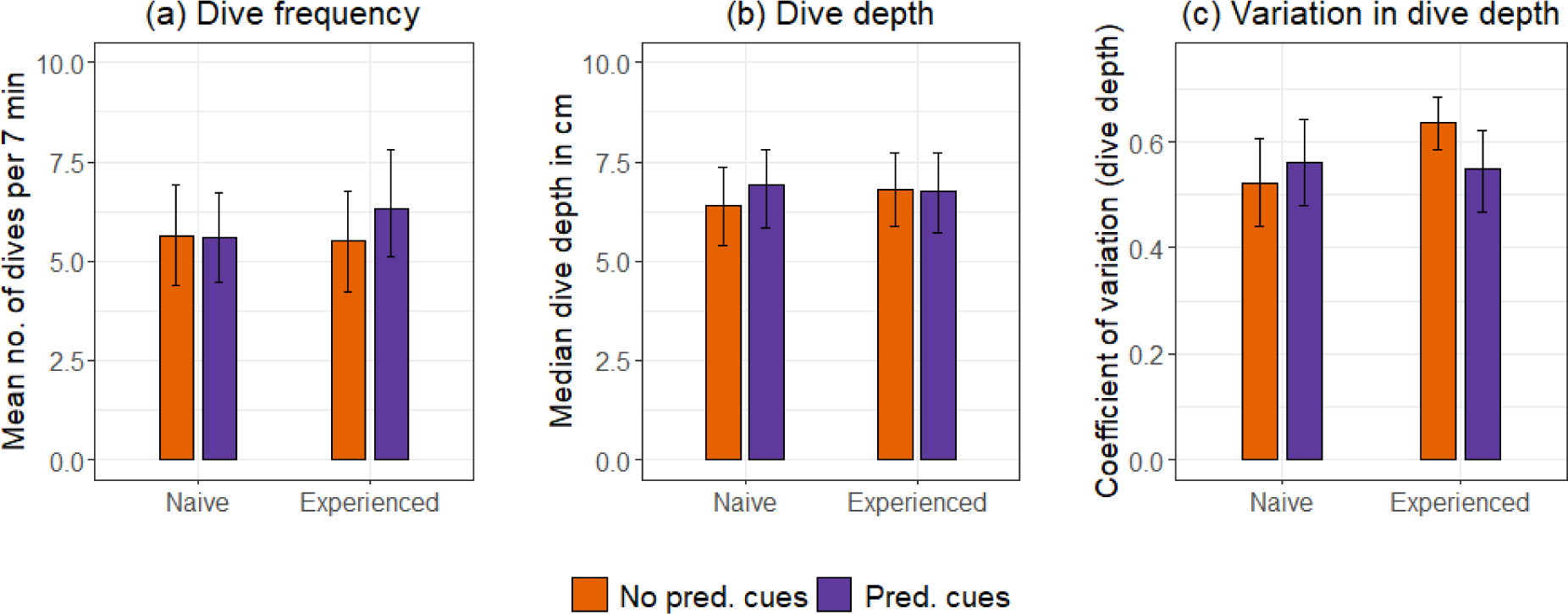
Comparison of (a) dive frequency, (b) median dive depth, and (c) coefficient of variation (CV) of dive depth between naive and experienced pupae at two different threat levels under normal resource-rich conditions. Orange represents the absence (no predation cues), and violet represents the presence of an immediate threat (predation cues). The means with 95% bootstrap confidence intervals are shown.

**Table 5:** Model estimates with 95% confidence intervals from three models: negative binomial GLM with dive frequency and linear models with median dive depth and CV dive depth as response variables. We estimated the effects of the larval growth environment (experienced and naive), behavioural assay environment (predation cue and control water) and their interaction on the response variables under resource-rich conditions. Entries in bold indicate that the effects are detectable.

|  | Dive frequency |  | Dive depth |  | Variation in dive depth |  |
| --- | --- | --- | --- | --- | --- | --- |
|  | Estimate/<br>coefficient | 95% Conf.<br>interval | Estimate/<br>coefficient | 95% Conf.<br>interval | Estimate/<br>coefficient | 95% Conf.<br>interval |
| Intercept<br>(block1:naive:control) | 1.819 | 1.557<br>2.088 | 6.562 | 5.444<br>7.680 | 0.471 | 0.383<br>0.559 |
| block2 | -0.351 | **-0.648**<br>**-0.052** | 0.411 | -0.873<br>1.696 | 0.070 | -0.036<br>0.177 |
| block3 | 0.021 | -0.234<br>0.279 | -0.819 | -1.933<br>0.293 | 0.099 | **0.012**<br>**0.186** |
| experienced | -0.058 | -0.380<br>0.263 | 0.430 | -0.971<br>1.832 | 0.112 | **0.000**<br>**0.224** |
| cue | -0.035 | -0.356<br>0.285 | 0.499 | -0.857<br>1.855 | 0.038 | -0.070<br>0.147 |
| experienced:cue | 0.173 | -0.278<br>0.625 | -0.525 | -2.481<br>1.431 | -0.121 | -0.278<br>0.034 |

**Table 6:** Likelihood ratio tests and F tests for the response variables– dive frequency, median, and CV dive depth under resource-rich conditions. We first assessed the interaction term by comparing models with the interaction term, ‘Larval growth env:Behavioural assay env’, and the nested simple models (without the interaction term). Models with the interaction term did not explain the data better than the simpler models. We used the models without the interaction term to assess the statistical significance of the main effects (larval growth environment, behavioural assay environment) through likelihood ratio tests and F tests. We used the ‘drop1’ command to perform the likelihood ratio tests for dive frequency models and the F tests for dive depth and variation in dive depth models.

|  | Dive frequency |  |  | Dive depth |  |  | Variation in dive depth |  |  |
| --- | --- | --- | --- | --- | --- | --- | --- | --- | --- |
| Deleted parameter | Likelihood ratio (chisq) | Degrees of freedom | P value | F value | Degrees of freedom | P value | F value | Degrees of freedom | P value |
| Larval growth env: Beh assay env | 0.566 | 1 | 0.451 | 0.281 | 1 | 0.596 | 2.379 | 1 | 0.125 |
| Block | 6.370 | 2 | 0.041 | 1.871 | 2 | 0.157 | 2.725 | 2 | 0.069 |
| Larval growth env | 0.062 | 1 | 0.802 | 0.107 | 1 | 0.744 | 1.566 | 1 | 0.212 |
| Beh assay env | 0.202 | 1 | 0.652 | 0.250 | 1 | 0.617 | 0.288 | 1 | 0.591 |

### Space use and Activity (Resource-rich conditions)

Space use and activity patterns did not change based on larval experience and the immediate threat environment under resource-rich conditions (Table 7 & Table 8). Pupae overall spent more time at the surface (60%) than middle column (27%) and bottom (13%). They also spent roughly 40% of their time active and 60% passive (Fig. 5).

**Fig. 5:**
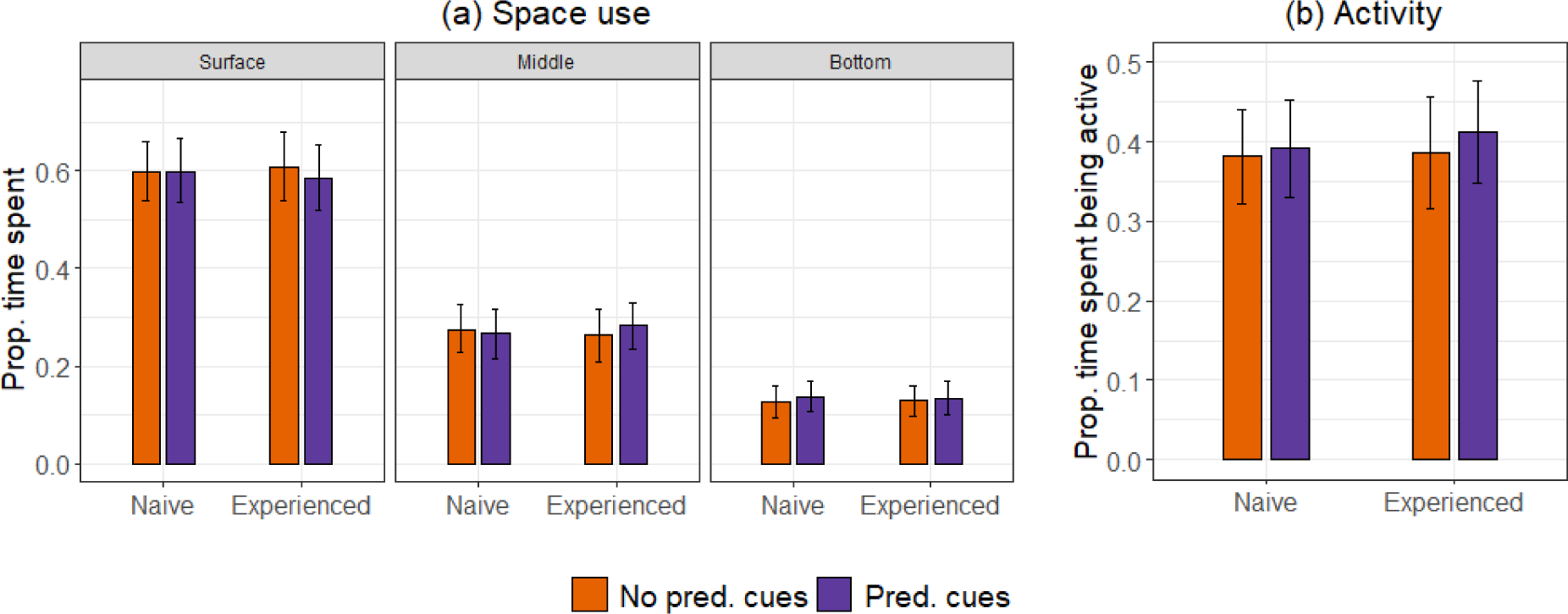
(a) The plots compare the proportion of time spent at the surface, middle and bottom layers by naive and experienced pupae across two threat levels (no predation cues and predation cues) under resource-rich conditions. (b) Comparison of the proportion of time spent being active by experienced and naive pupae across two threat levels. The means with 95% bootstrap confidence intervals are shown.

**Table 7:** Model estimates with 95% confidence intervals from two models: a multinomial model with proportions of time spent at the surface and bottom layers relative to the middle layer as response variables; a beta regression GLM with ‘proportion time spent being active’ as the response variable. We estimated the effects of the larval growth environment (experienced and naive), behavioural assay environment (predation cue and control water), and their interaction on the response variables under resource-rich conditions. Entries in bold indicate that the effects are detectable.

|  | Space use:<br>Surface/Middle column |  | Space use:<br>Bottom layer/Middle column |  | Activity |  |
| --- | --- | --- | --- | --- | --- | --- |
|  | Estimate/<br>coefficient | 95% Conf.<br>interval | Estimate/<br>coefficient | 95% Conf.<br>interval | Estimate/<br>coefficient | 95% Conf.<br>interval |
| Intercept<br>(block1:naive:con) | 0.617 | 0.331<br>0.903 | -0.904 | -1.326<br>-0.483 | -0.462 | -0.820<br>-0.104 |
| block2 | 0.732 | **0.386**<br>**1.077** | 0.288 | -0.222<br>0.799 | -0.706 | -1.107<br>-0.305 |
| block3 | 0.003 | -0.281<br>0.287 | 0.211 | -0.192<br>0.614 | -0.061 | -0.416<br>0.294 |
| experienced | 0.049 | -0.309<br>0.407 | 0.070 | -0.450<br>0.590 | -0.148 | -0.587<br>0.289 |
| cue | 0.034 | -0.323<br>0.393 | 0.121 | -0.394<br>0.637 | 0.140 | -0.296<br>0.577 |
| experienced:cue | -0.135 | -0.641<br>0.371 | -0.165 | -0.894<br>0.563 | 0.166 | -0.451<br>0.785 |

**Table 8:** Likelihood ratio tests for the response variables– proportion time spent at surface, middle, and bottom layers under resource-rich conditions; proportion time spent being active. We first assessed the interaction term by comparing models with the interaction term, ‘Larval growth env:Behavioural assay env’, and the nested simple models (without the interaction term). Models with the interaction term did not explain the data better than the simpler models. We used the models without the interaction term to assess the statistical significance of the main effects (larval growth environment, behavioural assay environment) through likelihood ratio tests. We used the ‘lrtest’ command from the package, ‘lmtest’, to perform the likelihood ratio tests for space use and activity models.

|  | Space use |  |  | Activity |  |  |
| --- | --- | --- | --- | --- | --- | --- |
| Deleted parameter | Likelihood ratio (chisq) | Degrees of freedom | P value | Likelihood ratio (chisq) | Degrees of freedom | P value |
| Larval growth env:Beh assay env | 1.777 | 2 | 0.411 | 0.279 | 1 | 0.597 |
| Block | 139.990 | 4 | <0.0001 | 13.383 | 2 | 0.001 |
| Larval growth env | 0.115 | 2 | 0.943 | 0.163 | 1 | 0.685 |
| Beh assay env | 1.112 | 2 | 0.573 | 2.033 | 1 | 0.153 |

## Discussion

Our study clearly demonstrates the behavioural carryover of predation risk across different life stages, highlighting the role of predation pressure in the evolution of complex life cycles. We observed behavioural carryover of larval experience during the pupal stage under typical resource-limited conditions. Different behavioural traits were tested, revealing drastic shifts in movements associated with escaping a threat.

Early larval conditions strongly influenced the diving behaviour of pupae. Experienced pupae changed their diving patterns. They dove more frequently, preferred shallower depths, and showed more variation in their dive depths than naive pupae. Such movements can make a prey’s location more unpredictable and help evade attacks by a potential predator (Humphries & Driver, 1970; Richardson et al., 2018). Early life experience is essential for acquiring information about predators and enhancing any innate strategy of evading a predator during an encounter (Griffin et al., 2000; Krama et al., 2023; Maloney & McLean, 1995). Prior experience can alert the prey of a potential threat, and cause it to change its behaviour even in the absence of any obvious threat in its environment, as we observed in our study. Such carryover effects are quite prevalent in many taxa. For example, the offspring of experienced scincid lizards show a higher tongue flick response than naive offspring in the absence of any immediate threat in their environment (Shine & Downes, 1999). Wild great tits that have already listened to predator vocalisations show reduced singing and increased alarm calls even in the absence of predator calls (Abbey-Lee et al., 2016). Larvae of fathead minnow that are exposed to predation cues as eggs swim shorter distances than naive larvae (Crowder & Ward, 2022).

We also observed a shift in the behaviour of experienced pupae under immediate threat. The change in behaviour shows that the experienced pupae could sense the predation cues. It signifies the retention of larval learning (Alloway, 1972; Blackiston et al., 2008), specifically in the predation context. The experienced pupae, however, behaved contrary to our expectations under immediate threat. One explanation for the unexpected behaviour of experienced pupae could be the use of different tactics to avoid encounters and to safeguard themselves once the predator is detected (Lima & Dill, 1990). Astonishingly, the resultant behaviour in the presence of immediate threat was similar to that of naive pupae, which did not change in response to threat levels. The similar diving behaviour of experienced and naive pupae under immediate threat might be because of an innate survival response. The innate response may also explain the observed changes in the space use patterns of both experienced and naive pupae in response to immediate threats. We can establish this by testing whether pupal dive and space use response is threat-sensitive against a gradient of predation cues irrespective of larval experience.

Our findings contribute to the broad understanding of the carryover effects of predation risk in complex life cycles (McCauley et al., 2011; Pechenik, 2006; Roux et al., 2021; Zhang et al., 2023). We highlight the importance of early predation experience in shaping the evolution of behavioural traits in a complex life cycle with extreme metamorphosis.

While we observed behavioural carryover as a shift in diving patterns, space use patterns did not change based on larval experience. This result might be a consequence of aerial respiration in pupae, for which they have to spend a reasonable amount of time at the surface. It could also be a result of our interval recording technique. We performed interval recording every 10 seconds and found a similar pattern of space use for experienced and naive pupae. However, from our dive frequency data, we know that the resting surface bouts of experienced pupae were frequently interrupted by dives. Therefore, we can deduce that despite the similar broad space use pattern for experienced and naive pupae, the time spent at the surface during one bout was less for the experienced than for the naive pupae.

We saw no reduction in time spent at the bottom by pupae despite predators being at the bottom of the behavioural assay tanks. This unaffected behaviour is likely because of the absence of local cues. The chemical cues were uniformly distributed in water, and the nymphs were enclosed in glass beakers, obstructing all the local disturbances. Over many behavioural recordings, we observed the predators trying to attack the pupae through the glass barrier. The caged attacks did not stop the pupae from going near the predator beakers, indicating that pupae need chemical and vibrational cues to identify predators.

Activity patterns did not change between experienced and naive pupae. The pupal activity was not altered by immediate threat as well. This finding was similar to that of Chandrasegaran et al. 2020: the wriggling behaviour of pupae did not change in response to predation threat from guppies. While recording the broad activity pattern, we observed a great diversity of movements that pupae perform in water. Categorising their activity further into different kinds of movements can give us a better understanding of the role of these various manoeuvres and how predation risk affects them individually.

Our results show how certain behavioural traits change due to prior experience and under immediate threat. Dive-related traits showed clear differences, while activity and space use did not. Prey species have multiple antipredator traits in their arsenal, and they use different traits and their combinations given the cost and benefit of their usage in the given ecological context (Caro et al., 2016; Kikuchi et al., 2023). For example, impala, wildebeest and zebra use different suites of traits at different intensities based on various predator traits (Palmer & Packer, 2021).

We found a general difference in the behaviour of pupae under resource-rich and resource-limited conditions (Fig. 3 & Fig. 5). Pupae spent more time at the surface and were less active under resource-rich conditions than in resource-limited conditions. We did not see any effects of larval predation risk experience and immediate threat on pupal behaviour under resource-rich conditions. Abundant food can increase the growth and development rate, which can help the prey either gain size and escape the predator gape (Brönmark & Miner, 1992; McCormick et al., 2019) or evade the habitat and metamorphose into another stage (Benard, 2004; Sergio et al., 2021). Therefore, there can be different ways of minimising risks under different contexts. Resource conditions can affect the ability to manage trade-offs between traits, such as development vs defence. Pupae might invest in development and leave the habitat faster in resource-rich versus resource-limited conditions.

## Conclusion

Despite the physiological and evolutionary constraints, behavioural carryover of predation risk can occur between distinct life stages. We have demonstrated this for the first time in a larva-pupa system where they share the habitat and threat. It suggests that the evolution of two distinct life stages is not decoupled if there is an adaptive advantage of an experience for both life stages. Our findings also highlight the importance of different contexts, here resource availability, to observe such behavioural carry-overs of predation risk. Different contexts can change the costs and benefits of certain traits and, hence, shift the associated trade-offs.

Our study led to some surprising findings-1. Frequent shallow dives in experienced pupae without immediate threat; 2. Reversal of that behaviour, rendering it similar to naive pupae behaviour in the presence of immediate threat, 3. Different effects of food conditions on the behavioural carryover. To understand them further, one can look at the adaptive value of these resulting traits.

We found behavioural carryover between two distinct stages in a multistage complex life cycle. It is imperative to examine how the effect of selection pressure on multiple stages accumulates or interacts to affect overall fitness.

## Acknowledgements

We thank the Indian Institute of Science for providing institutional and logistic facilities. We are grateful for all the help in colony maintenance and knowledge transfer from Kanchana Gaonkar, Manvi Sharma, Karthikeyan Chandrasegaran, and Akshay P. We would also like to thank our project assistants– Gokul Bhaskaran, Shubhada Shirish More and Navina Mable Francis, and a strong team of interns– Subhiksha M Bharadwaj, Aranya Dhibar, Ashmita Baruah, Reva T, Jitty Alin Jacob Hareendran K M, Nidhi Yadav, Ishika and Sukanya for supporting us during the experiments, troubleshooting, data extraction and data cross checking. We thank our friends for their invaluable assistance during the fieldwork. We appreciate the fellowship support from the Ministry of Education of India and the grant support from the Department of Science and Technology-Science and Engineering Research Board (DST-SERB) for the Scientific and Useful Profound Research Advancement (SUPRA) grant, SPR/2020/000384. We were also supported by the funds from DST-FIST [Fund for Improvement of S&T Infrastructure; sanction number SR/FST/LSII-025/2009(C)], DBT-IISc Partnership Program (Phase II; Department of Biotechnology, Ministry of Science and Technology and Indian Institute of Science, India), CSIR (Council of Scientific and Industrial Research) 37(1636)/14/EMR-II and Indian Institute of Science. We acknowledge the Institutional Animal Ethics Committee’s (constituted by the Indian Institute of Science, CAF/ Ethics/605/2018) approval for breeding mosquitoes in the lab and conducting experiments with live insects.

